# An Open Benchmark for Systems Vaccinology: Insights from the CMI-PB Challenges

**DOI:** 10.64898/2026.08.25.746820

**Authors:** Pramod Shinde, Lisa Willemsen, Jiyeun Lee, Shelby Orfield, Ziyuan Ren, Minori Aoki, Nicola Thrupp, Aanya Gupta, Cheng-Chang Wu, Liran Mao, Chuxuan Li, Yuhao Tan, Tram Anh Nguyen, Ning-Shan Chang, Philipp S. L. Schäfer, Jian Xing, Cemil Can Ali Marandi, Burhan Sabuwala, CMI-PB Challenge Contestants, Joaquin Reyna, Jeremy P Gygi, Brendan Ha, James A. Overton, Tal Einav, Jason A. Greenbaum, Leying Guan, Mari Kojima, Ferhat Ay, Barry Grant, Steven H. Kleinstein, Bjoern Peters

## Abstract

Systems vaccinology approaches have identified factors affecting vaccine responses in multiple studies, but the ability of computational models to generalize these findings to unseen data remains unclear. We established a community resource to create and compare models predicting *B. pertussis* booster vaccination responses and put such modeling approaches to the test. We compiled multi-modal experimental training data from three independent cohorts (n=117 individuals), and asked investigators to predict vaccine responses in a cohort of 54 newly recruited individuals using only their pre-booster vaccination data. We benchmarked a total of 107 computational models. Top-performing models were characterized by workflows that prioritized rigorous data preprocessing, robust imputation of missing data, and the use of multi-omics integration or non-linear machine learning. We identified pre-existing antigen-specific antibody titers and baseline monocyte frequencies as the most consistent predictors of post-vaccination immunity, highlighting the dominant role of individual immune setpoints. We established the resulting datasets and evaluation framework as a community resource to advance predictive immunology and facilitate personalized vaccination strategies.

## Introduction

Systems vaccinology aims to improve our understanding of the immune mechanisms involved in the cascade of events leading to vaccine responses and to develop models that enable the prediction of individual responses based on immune signatures. Such models, derived from integrative multi-modal experimental datasets, have shown promise in capturing complex biological interactions underlying vaccine responsiveness ^1^. However, the performance of these models on independent data is rarely evaluated. Initiatives such as the Critical Assessment of Structure Prediction (CASP) contest have provided invaluable data for researchers in the field of protein structure prediction by allowing them to test and compare model predictions in a quantitative fashion against previously unpublished data^2–6^. Such an initiative has been missing for systems vaccinology, where models are typically trained and tested on datasets identified in their original publication. Consequently, direct comparisons between different models are challenging due to differences in training and test data.

To address the need for standardized benchmarking and continuous methodological improvement in computational vaccinology, we organized the Computational Models in Immunity-Pertussis Booster (CMI-PB; www.cmi-pb.org) initiative. We followed a competition-driven approach that has been proven to accelerate methodological progress in related computational biology fields^2,4–7^. Rather than relying on retrospective datasets, we generated multi-modal experimental data specifically for the purpose of evaluating predictions on unseen individuals and evaluated model performance in a blinded manner. This design enabled a fully blinded evaluation of predictive performance, separating model development from outcome assessment. Unlike prior efforts in systems immunology, CMI-PB couples prospective data generation with predefined prediction tasks and standardized evaluation metrics.

We selected the immune response to *Bordetella pertussis* (the bacterial species causing whooping cough) as the biological system for this challenge due to 1) its public health relevance^8,9^ and 2) the unique ability to study how switching from the whole-cell pertussis (wP) vaccine introduced in the US in the 1940s to the less reactogenic acellular vaccine (aP) in the mid-1990s^8,9^ has impacted vaccine responses and how this switch can be linked to the observed waning of immunity. While aP vaccines protect from whooping cough equivalent to that of wP vaccines in clinical trials covering the initial period after vaccination, questions have been raised about the long-term durability^10^ and protection against transmission, contributing to the resurgence of pertussis^11^. Specifically, an increase in pertussis outbreaks has been reported in various countries that have switched from wP to aP vaccines^12,13^. As a result, several studies have explored waning immunity following aP vaccination and characterized distinct immune profiles between aP- and wP-primed individuals^14–17^. These investigations showed that long-lasting differences in T cell polarization, proliferation, and memory recall persist despite subsequent Tdap booster vaccination, yet how this difference in immune responses is maintained over time between individuals remains unclear^17^. Unlike influenza, whose immunological correlates and vaccine responses have been extensively profiled through large-scale systems vaccinology studies^18–21^, the molecular and cellular determinants of protection against pertussis are comparatively understudied^22^. Waning immunity after aP vaccination and variable responses to booster vaccinations underscore the need for a deeper mechanistic understanding of vaccine-induced protection. This establishes pertussis vaccines as a high value target for computational vaccinology, where standardized datasets and predictive benchmarks can accelerate discovery and guide rational vaccine design. Although centered on pertussis, the challenge framework is designed to be extensible to other vaccines and immune perturbations.

We organized three CMI-PB vaccine response prediction challenges between August 2021 and December 2024. The first was an internal dry-run challenge^23,24^, in which we established and stress-tested our strategies for clinical recruitment, experimental protocols, data handling, and selection and scoring of prediction tasks. We enrolled 96 individuals, 60 for training and 36 for challenge, from whom blood samples were drawn before and after vaccination. As an internal challenge, we asked members of our teams to predict selected readouts of post-vaccination antibody titers, cytokine production, cell frequencies, and gene expression given the pre-vaccination measurements. We were surprised to find that simple models performed well, often outperforming more complex models^23^. Specifically, chronological age (at time of booster) emerged as a competitive predictor of antigen-specific antibody titers, and pre-vaccination measurements of analytes targeted in a task showed high correlation with their post-vaccination measurements. We consequently defined two control models against which all future submissions would be benchmarked: *Control Model 1*, which ranks study individuals solely on their age (at time of booster), and *Control Model 2*, which ranks individuals by their pre-vaccination (day 0) levels of the analytes evaluated in each task. These controls ensure that computational models demonstrate true additional predictive value beyond readily available demographic or baseline data.

Following the dry-run, we launched an invited external challenge^25^ (n=117 training, n=21 challenge). In this phase, we also explicitly tested whether immune signatures derived from other vaccines (e.g., influenza, yellow fever) predict *B. pertussis* booster responses. We evaluated 22 published predictive models adapted from the systems immunology literature. None of these repurposed models outperformed the simple baseline control models when applied to the pertussis context^25^. These results highlighted limitations in current systems vaccinology: immune signatures appear highly context-specific and lack robust transferability across different vaccines^25^. On the other hand, the purpose-built models, trained specifically on CMI-PB data by invited contestants, achieved significant predictive success. The top-performing teams significantly outperformed both the age-based and baseline controls^25^. Winning architectures leveraged multi-omics integration (e.g., MCIA) and non-linear algorithms (e.g., CatBoost) to capture complex immune interactions, demonstrating that context-specific models can successfully decode the heterogeneity of vaccine responses^25^.

Building on these foundations^23,25^, we conducted a fully open (public) prediction challenge (**Figure 1**). Contestants trained models on standardized multi-modal data from 117 individuals and were evaluated on an independent cohort of 54 newly recruited individuals, providing an unbiased assessment of model generalizability. Here, we present the results of this public challenge and put them in the context of our broader experience building and refining this community resource for reproducible and benchmarked vaccine response prediction.

**Figure 1:**
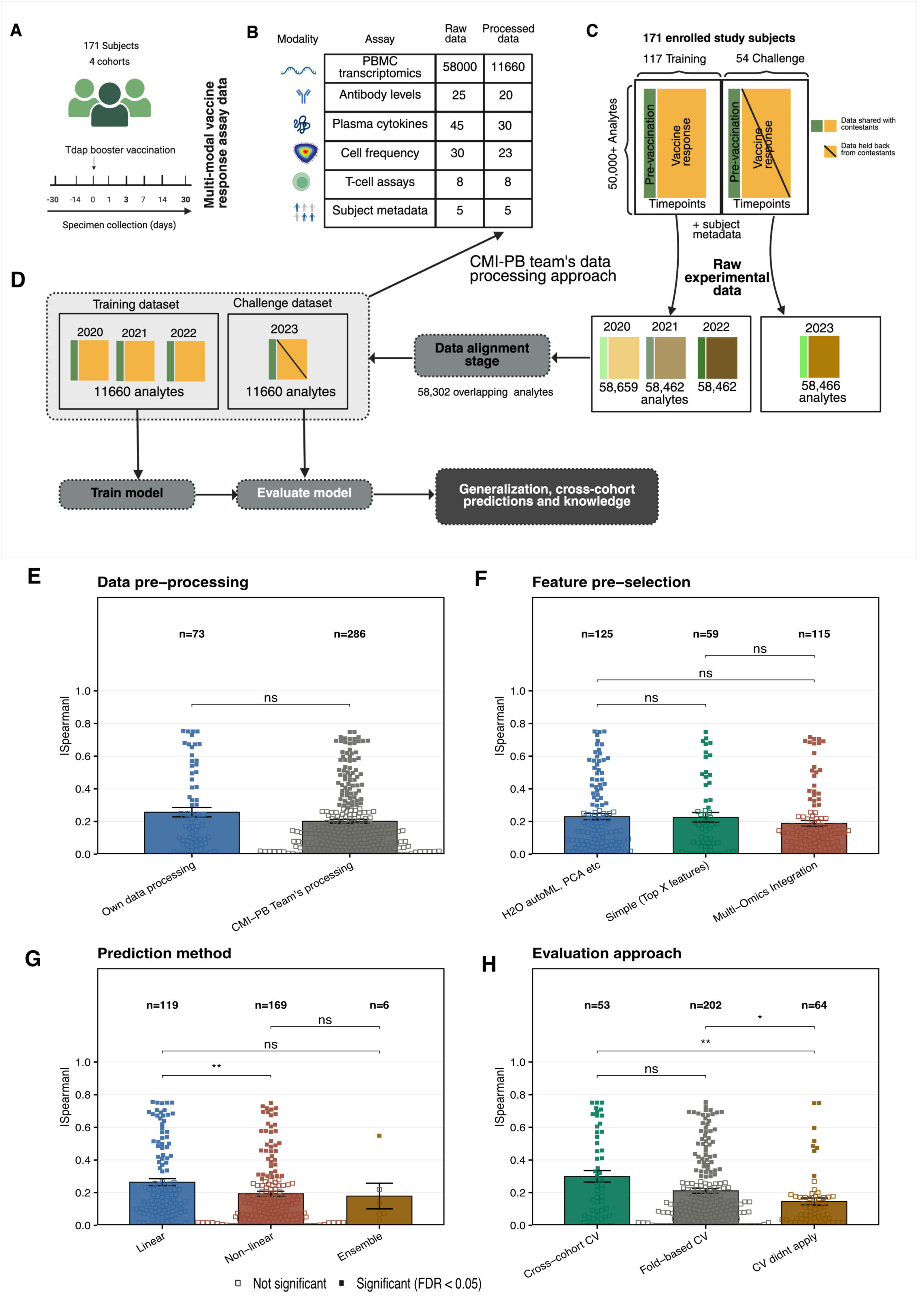
Overview of the multi-modal vaccine response data and modeling approaches. (A) Schematic of the study design for Tdap booster vaccination in 171 individuals across four cohorts, with biospecimen collection spanning pre- and post-vaccination timepoints. (B) Summary of multi-modal experimental assays performed, showing the number of analytes measured in raw and processed datasets for PBMC transcriptomics, antibody levels, plasma cytokines, cell frequencies, T-cell assays, and subject metadata. (C-D) Data processing and alignment pipeline integrating raw experimental data from four study years (2020-2023). From 58,000+ analytes measured per year, a shared set of 58,302 overlapping analytes was identified for downstream analysis. Data partitions illustrate how data from 2023 was withheld as a challenge dataset while previous years’ cohorts were used for training. (E-H) Performance of submitted models stratified by modeling design choices (mean |Spearman| ± SEM; filled squares: FDR < 0.05). (E-F) Data preprocessing source (E) and feature pre-selection strategy (F) had no significant impact on predictive performance. Linear models outperformed non-linear methods (G), and cross-cohort cross-validation was associated with higher performance compared to fold-based CV or no CV (H). Panels A, B, C were created with BioRender.com.

## Results

### Generating experimental data for training and testing prediction models

We generated assay data for samples from 54 newly recruited individuals. The assays covered four modalities that were also covered in previous CMI-PB challenges: i) bulk RNA sequencing of peripheral blood mononuclear cells (PBMCs), ii) plasma cytokine quantification, iii) PBMC subset frequency analysis, and iv) plasma antibody profiling against Tdap antigens. New to this challenge, we also captured T-cell activation and polarization assays (used in Task 4). Each study individual was followed for 28 days post-vaccination, with repeated blood samples collected both before (days -30, -14, and 0) and after (days 1, 3, 7, 14, and 28) the Tdap booster vaccination. Contestants were provided with a training dataset to build their prediction models that consisted of the 2020, 2021, and 2022 cohorts, for a total of 117 individuals (**Figure 1**)^23–25^, along with the baseline-only data for the 54 newly recruited individuals.

### Standardizing multi-modal data for cross-cohort modeling

Even under controlled conditions, experimental datasets generated at different times and by different operators are likely to exhibit systematic biases, which makes it hard to compare and transfer results from one dataset to another. To mitigate these effects, the CMI-PB team implemented a three-step preprocessing pipeline that involved 1) identification of common features across datasets, 2) baseline median normalization, and 3) batch-effect correction where those effects were apparent (detailed in **Methods and Figures S1, S2, and S3**). This standardized preprocessing was intended to reduce technical variability while preserving biologically meaningful signals across cohorts. As part of the preprocessing, the original ∼58,000 features were reduced to a subset of 11,660 robustly informative features: 11,589 transcriptomic, 28 cytokine, 20 antibody, and 23 PBMC cell subset features (**Figure 2B**). From the initial ∼57,500 transcriptomic features, we excluded mitochondrial genes, low-variance features, and lowly expressed genes (TPM < 1 in at least 30% of specimens). Challenge contestants were provided with both raw and processed data, processing scripts, and intermediate datasets, enabling them to adopt or modify specific preprocessing steps according to their modeling approaches.

**Figure 2.**
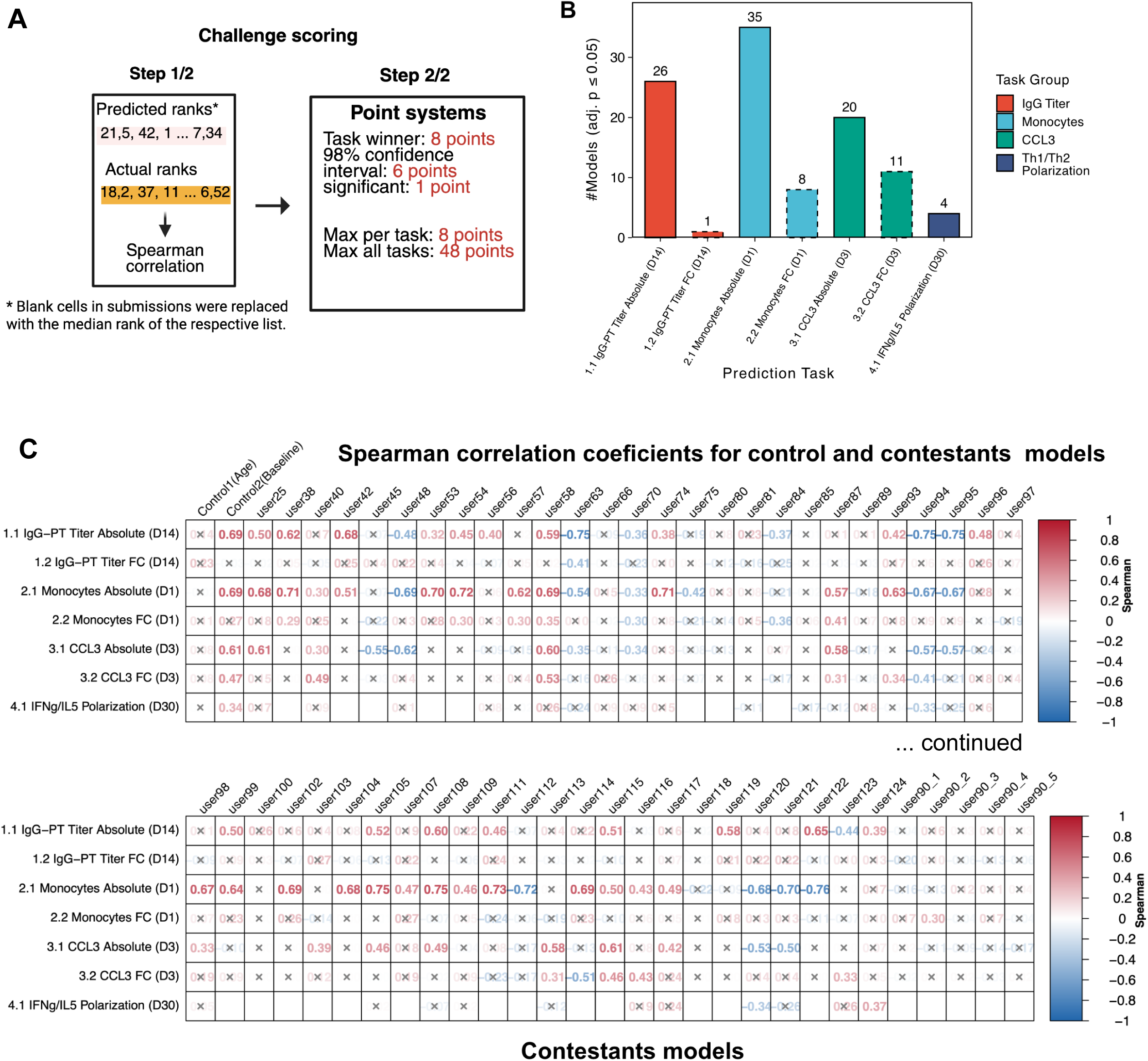
Evaluation framework and scoring of model submissions in the CMI-PB challenge. (A) Two-step scoring procedure for evaluating model submissions. In Step 1, predicted ranks submitted by contestants are compared to actual ranks of individuals based on measured immunological outcomes using Spearman correlation. In Step 2, points are awarded depending on performance. (B) Count of significant Spearman correlations for contestants models across the prediction tasks. (C) Heatmaps showing Spearman correlation coefficients for all contestant submissions across tasks. Each cell displays the correlation coefficient for a submission-task pair, with significant positive correlations highlighted in red and negative or near-zero correlations in blue. Gray “x” marks indicate missing or non-significant results. Submissions are arranged horizontally and tasks vertically to facilitate visual comparison of performance patterns across the challenge. Panel A is created with BioRender.com.

### Establishing prediction tasks that are appropriate for a public contest and that capture diverse aspects of the booster response

We learned from our previous challenges^23,25^ that when given a lot of tasks, many contestants only submitted results for a subset, and the feedback we received was that each new task came with substantial incremental effort. Thus, we wanted to limit the number of prediction tasks for the public contest while still covering the mechanistic diversity of vaccine responses we observed. We selected three biological readouts previously shown to be altered by booster vaccination in general and/or that were shown to differ between aP-primed and wP-primed individuals specifically^17,24^ as summarized in **Table 2**. This included the plasma IgG titers against pertussis toxin (PT) on day 14 post-booster vaccination (Task 1), PBMC monocyte frequencies on day 1 post-booster vaccination (Task 2), and *CCL3* gene expression in PBMCs on day 3 post-booster vaccination (Task 3). The *CCL3* transcript encodes a well-characterized proinflammatory chemokine that plays a critical role in the innate and early adaptive immune responses^24^. For each of these tasks, we asked for a ranking of the study individuals based on the absolute values (Task X.1) as well as a ranking of individuals based on the fold-change values compared to pre-vaccination (Task X.2). Additionally, a bonus prediction task focused on ranking individuals based on their IFN-γ/IL-5 cytokine production cell ratio at day 30 post-booster vaccination (Task 4.1), reflecting Th1/Th2 polarization, a known determinant of vaccine efficacy that differs between individuals primed with aP versus wP pertussis vaccines^17^. Given that the bonus task was newly introduced in this round, we kept its evaluation separate from the other Tasks.

**Table 1:** Overview of all CMI-PB prediction challenges. The CMI-PB project organized three consecutive prediction challenges to evaluate computational models predicting immune responses to *B. pertussis* booster vaccination based on multi-modal data. The first challenge was an internal “dry run” limited to consortium members and concluded in May 2022. The second “invited” challenge, completed in January 2024, included both consortium members and selected external researchers. The third “open” challenge, concluded in December 2024, invited participation from the broader public. Each challenge was built upon data and insights from previous rounds, with training datasets expanded by incorporating data from earlier challenges to improve the robustness and generalizability of predictions. Contestants submitted models to predict multiple immune readouts, and model performance was evaluated primarily using Spearman rank correlations between predicted and observed post-vaccination responses. We received 109 models across all three challenges.

| Prediction challenge title | Contestants | Study individuals per dataset (n) |  | Number of models received |
| --- | --- | --- | --- | --- |
|  |  | Training | Challenge |  |
| **First:** Internal dry run | CMI-PB consortium | 60 (28 aP + 32 wP) | 36 (19 aP + 17 wP) | 28 |
| **Second:** Invited challenge | Invited contestants | 96 (47 aP + 49 wP) | 21 (11 aP + 10 wP) | 25 |
| **Third:** Open Challenge | Public | 117 (58 aP + 59 wP) | 54 (27 aP + 27 wP) | 54 |

**Table 2.** Prediction tasks and model performance. Prediction tasks were grouped into antibody titer, cell frequency, and gene expression categories, each requiring contestants to rank individuals based on either absolute post-vaccination values or fold changes relative to baseline measurements. The table lists the task type, the specific immune readout, the number of submitted models for each task, how many models achieved statistically significant Spearman correlations, and the highest Spearman correlation observed. Absolute-value prediction tasks generally achieved stronger performance than fold-change tasks, with cell frequency predictions showing the highest overall correlations.

| Task ID | Task Type | Task statement | #models attempted tasks | #models with significant Spearman corr | Highest magnitude of Spearman correlation |
| --- | --- | --- | --- | --- | --- |
| 1.1 | Antigen-specific antibody titer | Rank the individuals by IgG antibody titers against PT that we detect in plasma 14 days post-booster vaccinations. | 54 | 24 (~45%) | -0.75 |
| 1.2 |  | Rank the individuals by fold change of IgG antibody titers against PT that we detect in plasma 14 days post-booster vaccinations, compared to their titer values at day 0. | 54 | 1 (~2%) | -0.41 |
| 2.1 | Cell subset frequency | Rank the individuals by predicted frequency of monocytes on day 1 post-booster vaccination. | 52 | 31 (~60%) | -0.76 |
| 2.2 |  | Rank the individuals by fold change of predicted frequency of monocytes on day 1 post-booster vaccination compared to their cell frequency values at day 0. | 51 | 5 (~10%) | 0.41 |
| 3.1 | Gene expression | Rank the individuals by predicted gene expression of *CCL3* on day 3 post-booster vaccination. | 41 | 19 (~46%) | -0.62 |
| 3.2 |  | Rank the individuals by fold change of predicted gene expression of *CCL3* on day 3 post-booster vaccination compared to *CCL3* gene expression values at day 0. | 42 | 9 (~21%) | 0.53 |
| 4.1<br>(Bonus task) | T-cell Polarization | Predict and rank individuals based on their Th1/Th2 (IFN-γ/IL-5) polarization ratio on day 30 post-booster vaccination. | 29 | 3 (~10%) | 0.37 |

### Evaluating the performance of control models on the public challenge dataset

We established two control models: 1) using individuals’ age (at the time of the booster) to rank them, and 2) using the pre-vaccination measurements of the analyte being evaluated to rank study individuals. Both control models performed well in the previous dry-run^23^ and invited challenges^25^, and their performance set a baseline that trained models should exceed. When testing the performance of Control Model 1 (age ranking), it failed to yield significant correlations for any of the six prediction tasks (**Figure 2C**). For the absolute-response tasks (Tasks X.1), the observed correlations were consistently positive but did not reach statistical significance. For the fold-change tasks (Tasks X.2), rankings were submitted in reverse order to reflect the expected inverse relationship with baseline values; however, these correlations also failed to reach significance, despite being directionally consistent with prior expectations. This was surprising to us but might be due to the limited age range of individuals (18 to 49 years) evaluated in this cohort.

In contrast, Control Model 2 (baseline ranking) proved highly predictive (**Figure 2C**). By simply ranking individuals based on their pre-vaccination levels of an analyte, this model achieved significant Spearman correlations for five of the seven tasks, including IgG-PT titers (Task 1.1), monocyte frequencies (Task 2.1), *CCL3* gene expression (Tasks 3.1, 3.2), and the pre-vaccination IFN-γ/IL-5 producing cells ratio (Task 4.1) (**Figure 2C**). As expected, baseline measurements were highly informative for absolute-response tasks, where positive correlations confirmed that individuals with higher pre-vaccination levels mounted stronger absolute responses (Tasks 1.1, 2.1, 3.1). For fold-change tasks where baseline was significant (Task 3.2), correlations were instead negative, consistent with regression to the mean, i.e., individuals starting lower having greater capacity for relative expansion. However, baseline levels failed to predict fold-change outcomes for tasks 1.2 and 2.2. This reflects a biological “ceiling effect,” where high pre-existing immunity limits the potential for further expansion^26^. Together, these results establish that pre-vaccination immune state is the dominant driver of post-vaccination responses across most tasks, while also revealing the limits of this signal, particularly for fold-change outcomes where inter-individual variability remains largely unexplained (**Figure S7**).

### Computational strategies employed by contestants in the public challenge

Overall, 54 computational models were submitted as part of the public challenge. Detailed methodological documentation and accessible code are available for models whose teams submitted their code, enabling comparisons across diverse modeling strategies (see **Supplementary Note S1**). Engagement was high across the tasks, with the majority of teams (48 out of 54) attempting all six prediction tasks. The methods employed spanned a broad spectrum of statistical and machine learning approaches, including linear regression, regression trees, partial least-squares (PLS), principal component regression, ensemble methods, and sparse regularization techniques.

### Employed data preprocessing strategies

Data preprocessing choices varied considerably across different teams. Most teams (40 out of 54) used the preprocessed datasets provided by us (the organizing team), which were curated to minimize batch effects and normalize data across multiple modalities (**Figure 1D**). Fourteen teams, including the first-place team (user58), opted to work directly with raw data, implementing their own data preprocessing workflows (**Figure 3B**). These teams often included established normalization procedures such as log₂(TPM + 1) transformation for gene expression, z-score standardization, and categorical encoding. In addition to the data transformation methods discussed above, teams working directly with raw datasets implemented a range of methods to handle missing data: while some teams used simple placeholder techniques (e.g., replacing missing values with NaNs), others used sophisticated algorithms such as k-nearest neighbors, SoftImpute, or tensor decomposition-based approaches. These advanced methods capitalized on correlations across samples, features, and timepoints to infer missing values with greater confidence.

**Figure 3.**
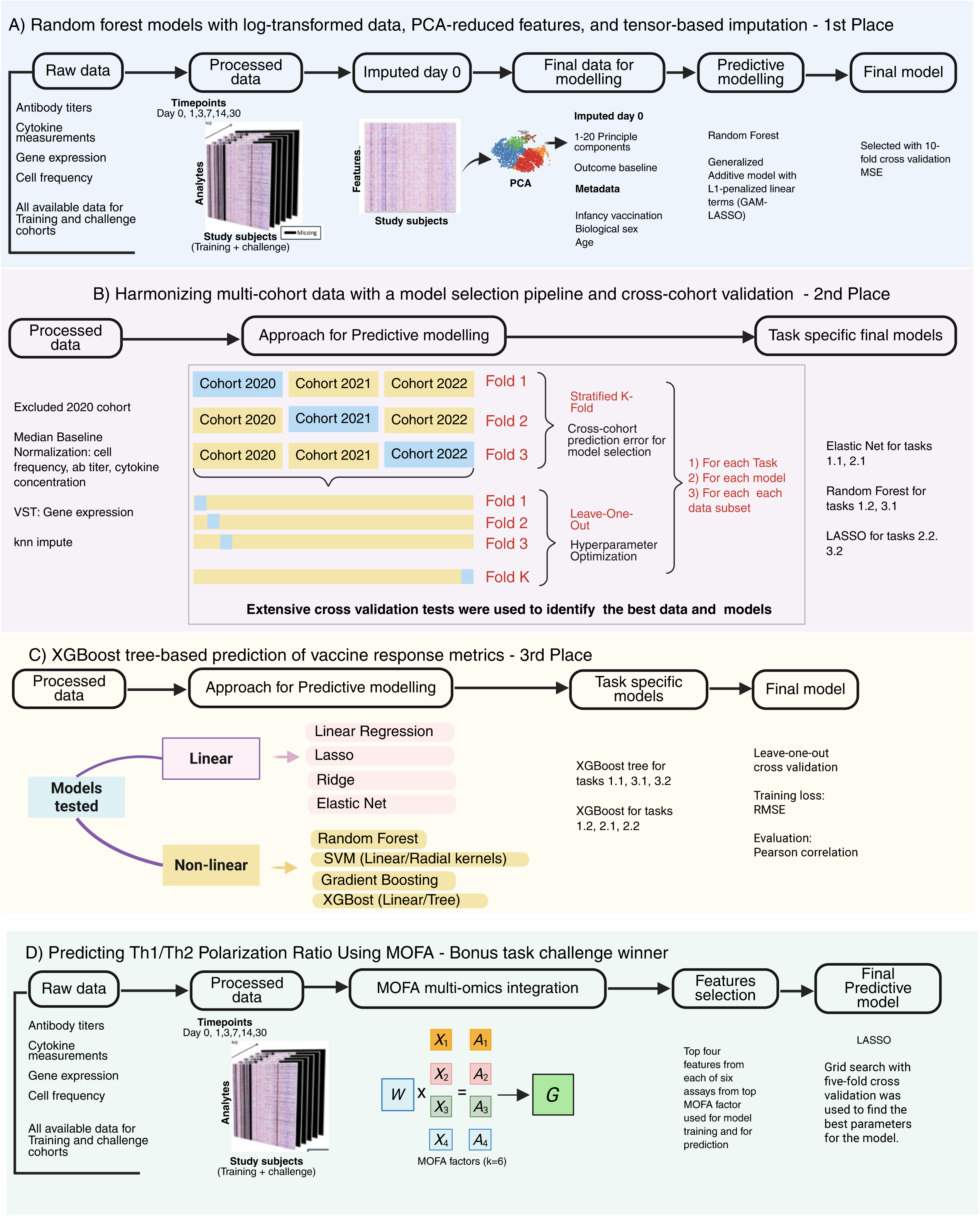
Summary of top-performing modeling strategies from the CMI-PB challenge. This figure illustrates the approaches used by winning teams across core and bonus tasks. (A) 1st Place: Random Forest models trained on PCA-reduced, log-transformed data with tensor-based imputation of day 0 values. Top predictors included baseline features and demographic metadata. Models were validated using 10-fold cross-validation. (B) 2nd Place: Cross-cohort modeling pipeline with stratified K-fold and leave-one-out validation. Normalization and feature selection were tailored by modality. Final models included elastic net, LASSO, and random forest. (C) 3rd Place: XGBoost tree-based models outperformed linear models for most tasks. Models were evaluated using leave-one-out cross-validation with RMSE and correlation metrics. (D) Bonus Task Winner: MOFA was used to integrate six omics layers. Latent factors were selected and refined using LASSO with 5-fold cross-validation to predict Th1/Th2 cytokine polarization. Created with BioRender.com.

### Employed modeling strategies

Several teams employed explicit multi-modal integration frameworks, including methods like Multi-Omics Factor Analysis (MOFA) or Joint and Individual Variation Explained (JIVE) (**Figure 1D**)^27,28^. While some teams utilized assay-specific features without reducing dimensionality, others adopted dimensionality reduction techniques such as Principal Component Analysis (PCA), factor analysis, or autoencoder-based embeddings to manage the high dimensionality inherent to multi-modal data. Others, like the bonus task-winning team from Stanford University (user124), integrated multi-modal assays while choosing to preserve feature resolution, opting out of dimensionality reduction to retain potentially informative variance across datasets.

Most submissions implemented supervised learning frameworks aligned with regression-style evaluation metrics. For example, the SPEAR approach (user42) utilizes a variational Bayesian framework to derive sparse latent factors from integrated multi-omics datasets, specifically weighting the model to optimize prediction of the response variable while maintaining interpretable feature signatures^29^. Teams explored both linear models (e.g., LASSO, ElasticNet, and Ridge) and non-linear models (e.g., XGBoost, Random Forest, CatBoost, and deep neural networks)^30,31^ (**Figure 1D**). Final model pipelines were often selected based on performance across K-fold or leave-one-out cross-validation schemes, a critical step for robust evaluation in small-sample, high-dimensional contexts^32^.

Feature selection was an integral component of many modeling pipelines, ranging from heuristic filtering and correlation-based ranking to algorithmic strategies such as regularization (LASSO), feature importance from tree-based models, or recursive feature elimination (**Figure 1D**). While several teams reported top predictors, such as specific immune cell subsets or cytokines, others operated in lower-dimensional latent spaces where interpretability was less direct.

### Shared strategies of top-performing predictions

Among the 54 models submitted, the three top-performing models differed significantly in their computational choices and yet converged on several methodological principles that likely contributed to their success. Each approach emphasized rigorous data preprocessing, particularly data normalization, standardization, and batch-effect correction, indicating that careful handling of technical variability across multiple omics assays was a prerequisite for better predictions (model details are presented in **Figure 3**). This convergence suggests that model performance was driven more by strategy than by any single algorithmic choice.

The first-place team, led by Cheng-Chang Wu from the University of Minnesota, utilized a random forest algorithm enhanced by PCA-derived features and tensor-based imputation. This pipeline specifically leveraged PCA to effectively reduce the dimensionality of high-dimensional multi-omics data and implemented tensor decomposition imputation to address missing data comprehensively across samples, features, and timepoints (**Figure 3A**). On the other hand, the second-place team, led by Philipp Schäfer from University Hospital Heidelberg, considered regularized regression models (ElasticNet and LASSO) and Random Forest approaches in a cohort-aware manner. Their method included selective dataset exclusion, stringent nested cross-validation (outer group k-fold and inner leave-one-out cross-validation), and systematic management of batch effects to maximize model robustness and interpretability (**Figure 3B**). The third-place model, led by Liran Mao from the University of Pennsylvania, employed XGBoost, a tree-based ensemble algorithm^33^, optimized by careful preprocessing, median imputation, and minimal dimensionality reduction, achieving particularly high performance for absolute immune response tasks (**Figure 3C**).

All top-performing models implemented tailored imputation strategies instead of strictly limiting analyses to complete cases. The first-place model’s tensor-based imputation exploited complex correlations across multiple dimensions. The second-place team selectively excluded problematic datasets rather than employing complex imputation, focusing instead on robust quality control. The third-place team applied straightforward median imputation, prioritizing the retention of biological detail while effectively handling sparsity.

To address overfitting, a common challenge in high-dimensional data modeling, the top-performing teams employed rigorous validation strategies. The first-place team used 10-fold cross-validation guided by mean squared error (MSE) to mitigate variance and prevent overfitting. The second-place team’s nested cross-validation minimized variance across test folds, reflecting robustness across multiple cohorts. The third-place team relied on leave-one-out cross-validation optimized on root mean squared error (RMSE) for hyperparameter tuning, ensuring stable predictions across data subsets. Together, these validation choices helped distinguish models that generalized well from those prone to overfitting.

Beyond architectural differences, all three methods included the baseline immune states , which are also utilized by Control Model 2, as dominant predictors of subsequent vaccine-induced responses (**Table 3**). The importance of baseline features is reflected in the performance: for absolute-response tasks, where the baseline is most informative, the top teams closely matched or exceeded control model 2. On Task 1.1 (IgG-PT absolute, D14), the 2nd-place team achieved the highest correlation overall (ρ = 0.751 vs. ρ = 0.694 for baseline), while the 1st- and 3rd-place teams fell slightly below (ρ = 0.587 and ρ = 0.114, respectively). On Task 2.1 (monocytes absolute, D1), all three teams tracked closely with baseline (ρ = 0.693), with 1st- and 3rd-placed teams matching it almost exactly (ρ = 0.695 and ρ = 0.572). Similarly, on Task 3.1 (CCL3 absolute, D3), 1st- and 3rd-placed teams approached *control model 2*’s performance (ρ = 0.598 and ρ = 0.578 vs. ρ = 0.609). For fold-change tasks, where baseline is a weaker predictor, the 2nd-place team was notably the only model to achieve a meaningful positive correlation on Task 1.2 (IgG-PT FC, ρ = 0.408), suggesting its modeling strategy captured variance beyond pre-existing immunity. Together, these results underscore that vaccine responses are substantially shaped by prior immune state, and the marginal gains of complex models over a simple baseline ranking were modest for most tasks.

### Extracting biological variables driving prediction performance

Identifying the biological features that a model uses for successful predictions is not trivial for the non-mechanistic models employed by most contestants, as there is no straightforward link from input features to prediction outcomes. This is further complicated by the broad range of modeling strategies employed, each of which would have to be examined separately. In addition, feature importance is not uniquely defined and can be quantified in multiple ways depending on the modeling approach. For example, in random forest models, feature relevance can be assessed using permutation feature importance (a model-agnostic approach that measures the drop in predictive performance after permuting a feature) or by mean decrease in impurity, defined as the average reduction in mean squared error (for regression) or Gini impurity/entropy (for classification) across splits that use that feature.

To enable at least a qualitative cross-model comparison, we asked contestants to submit the top features their models relied on. We then compiled a comprehensive list of informative features from all models that scored a significant correlation coefficient for specific tasks. “Informative features” refer to both observed variables, such as individual assay or demographic features (e.g., antibody titers or age at booster vaccination), and unobserved variables that capture and summarize patterns in the data, such as components (principal component analysis), latent variables or factors (factor analysis), or embeddings (deep learning models).

### Biological variables predicting day 14 post-vaccination IgG-PT responses (tasks 1.X)

For Task 1.1, which asked to predict absolute IgG-PT titers on day 14 post-vaccination, 25 of the 54 submitted models demonstrated statistically significant Spearman correlations (**Table 2**). Of the models that disclosed their informative feature set (15 of 19), the most consistently selected predictor was the baseline IgG-PT titers, present in 10 of these 15 models, including the 1st-place team (user63) (**Figure 4B**). The recurrence of baseline antibody titers as predictors underscores the importance of pre-existing humoral immunity as a primary determinant of post-booster antibody responses. This likely indicates the presence and functionality of PT-specific memory B cells, formed after earlier vaccinations or natural exposures, facilitating antibody responses upon subsequent vaccine PT exposure^34^.

**Figure 4.**
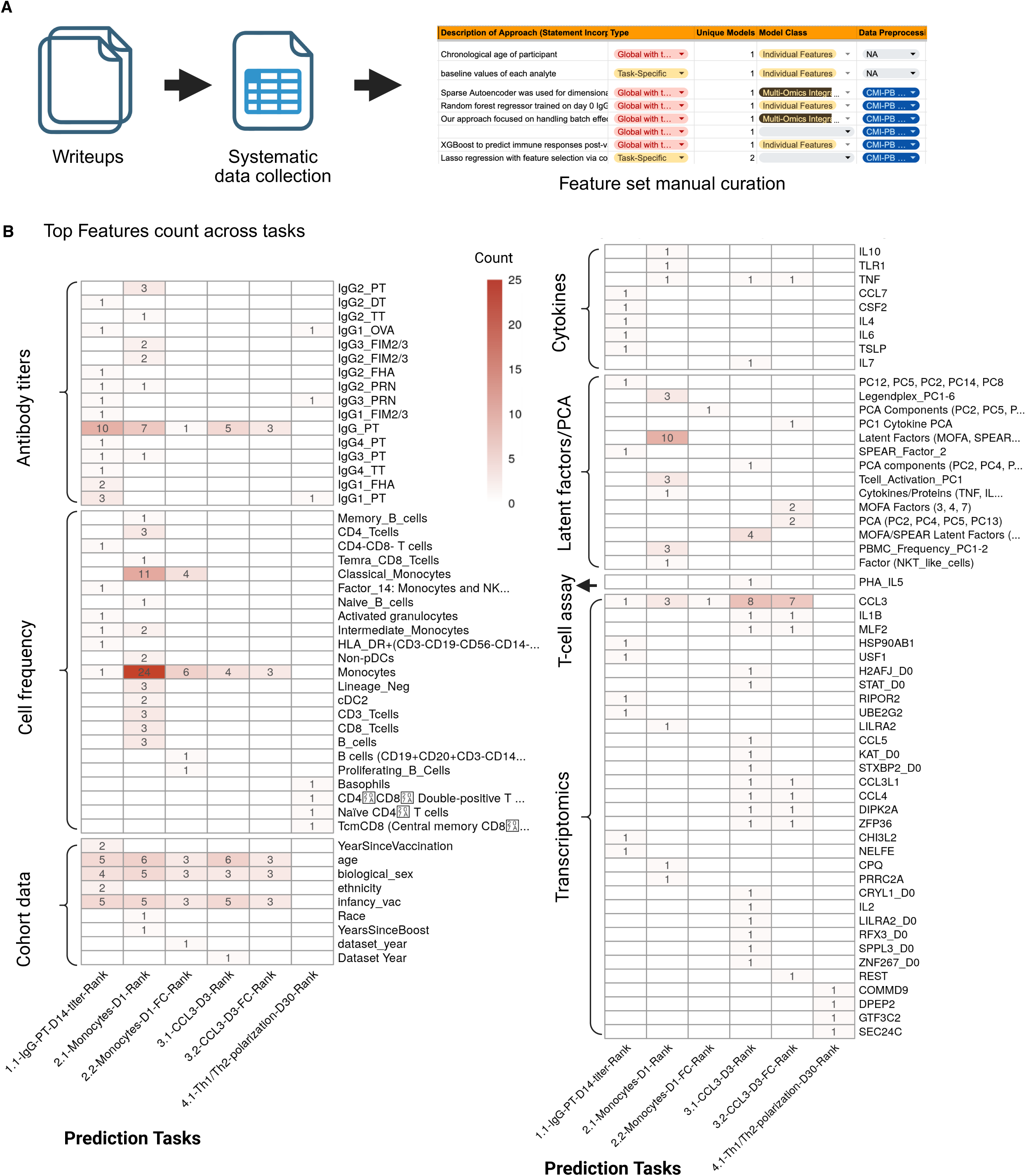
The most important features identified by models showed significant correlation coefficients per task. A) Free-text method write-ups from all teams were manually curated into a structured table recording features used, model type (global or task-specific), model class, and preprocessing and standardized them. B) Number of models naming each feature as a top contributor, per task. Rows are features grouped by assay; columns are the six tasks. Colour and printed number give the count; empty cells mean the feature was not reported. Counts include only models with a significant Spearman correlation for that task (P < 0.05). Feature names are as reported by teams; latent factors and principal components are model-specific and not comparable across submissions.

Given the dominant impact of pre-existing antibody titers on post-vaccination antibody titers, other biological factors should be more apparent when evaluating Task 1.2, where the fold change in antibody titers on day 14 compared to baseline is evaluated. Out of 54 submitted models, only one model (user63) achieved a significant Spearman correlation coefficient (rho = -0.41). The selected feature from this model was baseline monocyte frequencies, which was also one of key features listed in Task 1.1. Other features included the percentages of CD4⁺ T cells and T-cell polarization (IFN-γ/IL-5 producing cells) after PHA and PT stimulation. In summary, predictions of vaccine-induced IgG-PT antibody responses were largely driven by pre-vaccination IgG-PT, monocyte and CD4+ T cell frequency, and CD4+ T cell polarization impacting the relative change in antibody titers.

### Biological variables predicting day 1 post-vaccination monocyte frequency (tasks 2.X)

Of the submitted predictions for Task 2.1, 26 out of 54 models had a significant Spearman correlation coefficient, and 24 of the models had submitted a list of predictors. The most frequently selected feature was baseline monocyte frequency (Monocytes_D0), appearing in 24 of these models. Similar to Task 1.1, this underlines that a great starting point for predicting the absolute value of monocyte frequencies post-booster vaccination is to establish the host’s innate baseline prior to vaccination.

Task 2.2 asked for the relative change in monocyte frequencies on day 1. Out of 51 submitted models, six models achieved a significant Spearman correlation coefficient, and we received a list of predictors from five models (**Figure 4B**). Of these five models, four identified baseline monocyte frequencies (Monocytes_D0) as the key predictor. This again highlights the robust selection of baseline monocyte frequencies, selected by 4 models, confirming their importance in defining subsequent changes in monocyte numbers. Classical monocyte subsets at baseline and day 1 (Classical_Monocytes_D0 and Classical_Monocytes_D1) further supported the notion that monocyte subset-specific composition may influence their dynamic response to vaccination. Of note, one model included proliferating B cells and antigen-experienced B cell subsets, suggesting a link between humoral adaptive responses and the magnitude of early innate monocyte responses.

Taken together, results from Task 2.1 and Task 2.2 highlight baseline circulating monocytes as primary determinants of monocyte frequency and their dynamic responses to booster vaccination. The integration of humoral, adaptive cellular subsets, and systemic cytokine profiles further emphasizes the integrated regulation of innate immune cell responses following Tdap booster immunization.

### Biological variables predicting day 3 post-vaccination *CCL3* gene expression (tasks 3.X)

For Task 3.1, 41 teams submitted their models, of which 19 models showed a significant Spearman correlation coefficient. Biological variables were compiled from 14 submitted models. The most frequently selected variable was baseline *CCL3* gene expression (CCL3_D0), present in 8 models (**Figure 4B**). The top-performing model for Task 3.1 (model48) included CCL3_D0, demographic variables, and baseline features but moderately outperformed the baseline-only control model, suggesting that while *CCL3* is predictable solely by its baseline, the predictions can be improved by including demographic variables and other baseline features.

Baseline monocyte frequency (Monocytes_D0) was selected in 4 models, supporting the contribution of myeloid cell setpoints to early inflammatory cytokine responses^35^. Latent factors derived from MOFA^36^, SPEAR^29^, or PCA were used in 4 models, indicating that multi-omic integration captures relevant underlying immune activation states. These latent factor representations likely summarize shared variation across cell frequencies, transcriptomics, and cytokine activities, enabling models to leverage immune signals from multiple modalities that are more predictive than individual measurements alone. A diverse array of gene expression features, including *RFX3*, *ZNF267*, *IL1B*, *TNF*, and *ZFP36*, were also included across models, reflecting potential transcriptional regulators or downstream products of innate signaling cascades. *IL1B* and *TNF* encode pro-inflammatory cytokines central to innate immune activation, particularly in monocytes and macrophages. ZFP36 regulates the stability of inflammatory mRNAs, including *TNF* transcripts, while *RFX3* and *ZNF267* are transcriptional regulators that may modulate broader immune gene expression programs.

For Task 3.2, focused on predicting *CCL3* fold-change (day 3 vs. day 0), 42 teams submitted their models, of which 9 models showed significant Spearman correlation coefficients, and predictors were compiled from 8 models (**Figure 4B**). CCL3_D0 again emerged as the dominant feature, selected in 7 models, followed by demographic features (age, biological sex, and infancy vaccination; 3 models each) and Monocytes_D0 (3 models). This indicates that both absolute and relative *CCL3* responses are rooted in pre-existing innate setpoints and demographic context. IgG_PT_D0 was also selected in 3 models, highlighting a potential link between humoral immune setpoints and inflammatory regulation.

Two models incorporated MOFA latent factors and PCA components, suggesting shared molecular programs captured across multi-omic space. These latent variables may reflect coordinated activation of chemokine and interferon pathways. Other selected features included cytokine PCA axes and inflammatory gene expression markers such as *IL1B, CCL4, CCL3L1, DIPK2A*, and *ZFP36*, suggesting specific regulatory nodes that shape *CCL3* amplitude in response to Tdap.

Taken together, these findings indicate that *CCL3* responses following booster vaccination are strongly influenced by baseline gene expression levels, demographic factors, and integrated innate inflammatory states.

### Biological variables predicting day 30 post-vaccination Th1 polarization (Task 4)

For the bonus task, 29 teams attempted to rank individuals based on their IFN-γ/IL-5 polarization ratio on day 30 post-booster vaccination. Three teams (user124, user120, and user94), along with the baseline control model 1 (rho = 0.34), achieved a statistically significant Spearman correlation coefficient. The models from user124 (rho = 0.37) employed Multi-Omics Factor Analysis (MOFA) integration^36^, user94 (rho = 0.33) applied a regularized regression model with baseline features, and user120 (rho = 0.34) incorporated the IFN-γ/IL-5 polarization (same as control model 2) as part of their model building. All significant models identified baseline IFN-γ/IL-5 polarization as a primary predictor, emphasizing that pre-existing Th1/Th2 balance shapes the subsequent vaccine-induced trajectory.

The top-performing team, led by Aanya Gupta from Stanford University (user124), utilized MOFA to integrate assays from training (2020-2022) datasets in an unsupervised fashion^36^. Individual metadata features, including infancy vaccination type and biological sex, were encoded and included during model construction. The MOFA framework was configured to infer six latent factors under slow-convergence settings with sparsity-inducing regularization, yielding biologically interpretable summaries of shared variation across modalities. Their model inferred six biologically condensed factors using slow convergence and specific regularization (spike-and-slab weights) (described in **Figure 3D**). From each factor, the four highest-weight features were selected and used as inputs to a downstream elastic net regression model.

Hyperparameters were optimized using five-fold cross-validation with Spearman correlation as the objective metric. The final trained model was applied to the 2023 challenge cohort using an identical preprocessing pipeline, and ranked predictions were submitted for evaluation.

Key biological features identified across the successful models reflect the underlying complexity of immune regulation contributing to Th1/Th2 balance. The models incorporated diverse cellular subsets, including the percentage of basophils, which are potent inducers of Th2 polarization, alongside central memory CD8+ T cells associated with sustained Th1-type memory and CD4+CD8+ double-positive T cells implicated in multifunctional immune regulation.

Transcriptomic analysis highlighted specific genes such as *DPEP2, SEC24C, GTF3C2*, and *COMMD9* as potential proxies for regulatory pathways involved in immune activation. *SEC24C* and *GTF3C2* may reflect broader cellular activation and transcriptional activity. Furthermore, the inclusion of antigen-specific antibodies (including IgG1_PT, IgG1_OVA, and IgG3_PRN) captured the influence of prior antigen-specific humoral immunity and cumulative antigen exposure on subsequent polarization. While demographic factors like age were integrated into some models, such as that of user120, they did not show an independent association with the IFN-γ / IL-5 ratio, aligning with previous pertussis vaccine cohort studies^37^.

## Discussion

The results of our third CMI-PB Challenge provide a number of insights into what it takes to predict vaccine responses. Despite methodological diversity among winning teams, the 1st-, 2nd-, and 3rd-place pipelines converged on several modeling principles essential for extracting the signal from the CMI-PB dataset. First, aggressive data hygiene preceded modeling; all three teams devoted substantial effort to batch correction, variance-stabilizing transforms (log or VST), and feature filtering. This convergence suggests that preprocessing decisions are not merely technical details but are of importance for extensive data processing for downstream biological inference. Second, missing-value strategies were integral modeling steps rather than preprocessing afterthoughts. Whether via tensor decomposition (1st place), cohort-aware exclusion with conservative QC (2nd place), or median imputation (3rd place), each group demonstrated that handling missing data significantly shapes downstream performance. Third, winning models embedded biological knowledge, including infancy priming (aP vs. wP), sex, age, and baseline readouts, reflecting consensus on anchoring the prediction space in domain-specific context. Fourth, validation rigor to avoid overfitting by utilizing nested or leave-one-out cross-validation loops, emphasizing careful resampling over elaborate architecture tuning. Finally, the pipelines spanned a methodological spectrum, from feature-compressed ensembles (tensor-imputed PCs + Random Forest) and cohort-aware regularized regressions to high-capacity XGBoost models^33^ trained on minimally reduced data, suggesting that multiple algorithmic approaches can be successful. These results are in line with findings from our previous two challenges^23,25^, and this emphasizes that prediction success seems to hinge less on algorithm choice and more on disciplined workflow: meticulous preprocessing, principled handling of sparsity, inclusion of all relevant information, and robust cross-validation to avoid overfitting.

We split our tasks into two subtasks: the X.1) subtasks evaluate the ranking of individuals based on absolute readouts, while the X.2) Subtasks evaluate the ranking of individuals based on their fold change from pre- to post-vaccination. Thus, the performance on the X.1) subtasks was possible to predict reasonably well based on pre-existing responses alone, while the X.2) subtasks were not and, accordingly, showed much fewer models that succeeded. The comparatively weaker performance on fold-change tasks suggests that baseline measurements alone are insufficient to predict how much an individual’s immune response will increase after booster vaccination.

What do these prediction models tell us about the biological factors impacting responses to pertussis booster vaccination? The clearest finding is that the post-vaccination immune response is strongly anchored in the pre-vaccination state. Pre-vaccination readouts for antibody titers, cytokine production, cell type frequencies, and T cell polarization were all predictive of their respective post-vaccination values, as reflected in the strong performance of Control Model 2 across five of seven tasks (ρ = 0.47-0.69; **Figure 2C**, **Figure S7**). However, the pre-vaccination state alone did not fully account for inter-individual variability. For fold-change tasks, the baseline was a strong but not uniquely dominant predictor. On Tasks 1.1 and 2.1, several individual analytes outperformed baseline, indicating that additional immune features carry a meaningful predictive signal beyond the task-specific pre-vaccination measurement. On Task 3.1, baseline ranked among the top performers, suggesting that pre-vaccination CCL3 expression captures most of the available predictive signal for this readout. For fold-change tasks, Task 1.2 was largely unpredictable by any approach, with a single model reaching significance - consistent with the ceiling effect discussed above. Tasks 2.2 and 3.2 were more tractable, with a small number of models reaching significance despite their baselines performing modestly, suggesting that fold-change in these readouts is partially driven by features orthogonal to pre-existing levels. Baseline captures much of the rank structure but leaves a meaningful portion of the rank order unexplained. Complex models extracted additional signal from other immune features, and this distinction matters: we are not in a regime where “baseline is all you need” but rather one where baseline is the dominant signal and a substantial portion of the variance remains unexplained.

The features selected by top-performing models offer insight into what drives residual variability beyond the pre-vaccination baseline. Across fold-change tasks in particular, winning models consistently drew on both innate and adaptive immune compartments. Monocyte frequencies, alongside cytokines such as IL-6 and CSF2, were strong predictors of post-vaccination antibody titers. Adaptive immune features, including memory CD8⁺ T cells, CD4⁺CD8⁺ T cells, and cytokine secretion profiles such as IL-5 and IFN-γ, informed models predicting Th1/Th2 polarization. These findings suggest that effective vaccine response prediction must account for crosstalk between innate activation states and adaptive effector functions. More broadly, these modeling results are consistent with mechanistic studies showing that early IFN-γ production after booster vaccination is associated with the maintenance of Th1/Th2 polarization and that pre-existing memory T cells contribute to sustaining these polarization states^17^. The modeling framework and experimental studies thus provide complementary perspectives on the same underlying biological processes rather than redundant ones.

This pattern is not unique to pertussis - across vaccines, pre-vaccination immune state is consistently the dominant predictor of post-vaccination responses yet explains only a fraction of inter-individual variability^38,39^. For influenza, post-vaccination HAI titer prediction requires integrating titers against multiple historical antigenic variants alongside transcriptional and cellular baseline features, with pre-vaccination HAI against the vaccine strain alone performing poorly as a standalone predictor^38^. For individuals with lower baseline titers in particular, pre-vaccination gene expression provides independent explanatory power for residual response variation that titer-based models cannot capture^40^. More broadly, analysis of blood transcriptional profiles across 13 vaccines identified distinct pre-vaccination endotypes predictive of antibody responses, yet the generalizability and mechanisms underlying these associations remain incompletely defined^39^. The pertussis booster setting appears analogous: baseline immune state strongly constrains the likely rank-order response, but the factors driving residual inter-individual variability remain an open question.

This has wide-reaching implications. First of all, it impacts how we quantify vaccine responses and what we consider an effective response. Is an individual with an already high antibody titer that does not increase post-vaccination a ‘poor responder’? Is an individual who goes from a very low titer to a low titer a great responder? These scenarios illustrate that absolute magnitude and relative change reflect different biological properties and should not be interpreted interchangeably. This is further compounded by ceiling effects: individuals with high baseline responses have limited dynamic range for further increases, constraining fold-change independently of any true difference in vaccine-inducible immune capacity. Ideally, we would ask models to make quantitative predictions of biological variables, which implicitly allow us to define rankings between individuals and define changes between time points.

Ideally we would evaluate readouts like antibody titers based on their relevance to protection from disease, but such correlates have not been established for *B. pertussis*. We therefore structured the prediction tasks around reproducible immunological measurements such as antibody levels, cell frequencies, cytokines, and gene expression that allow consistent cross-cohort comparison and model evaluation.

Our studies established a comprehensive dataset and evaluation framework as an open community resource, which will be continuously available. We invite and encourage other scientists to analyze it and use it to examine their own hypotheses. By providing both raw and harmonized data alongside transparent evaluation metrics, we aim to promote reproducibility and iterative methodological improvement within the systems vaccinology community. While this third challenge completes the funding for our work on the pertussis booster vaccination grant, we continue and advance on this work for seasonal influenza vaccination (cmi-x.org), where we can benefit from the lessons learned in this context, and we invite the community to participate.

## Methods

### Ethics statement

This study was performed with approvals from the Institutional Review Board at the La Jolla Institute for Immunology, and written informed consent was obtained from all study individuals before enrollment (protocol number VD-101).

### Challenge data and ground truth

The public CMI-PB prediction challenge is outlined in **Figure 1**. A total of three multi-modal datasets were provided to contestants, consisting of 171 individuals. The entire dataset was split into training and challenge datasets. The training dataset includes two independent cohorts, the 2020, 2021, and 2022 cohorts, and these cohorts are described in detail in three recent publications: da Silva Antunes et al.^24^, Shinde et al. (first challenge)^23^, and Shinde et al. (second challenge)^25^. The challenge or ground truth evaluation dataset consists of 54 individuals, and we conducted experimental assays similar to those performed on the training datasets, as described in the following:

#### Experimental model and cohort details

The characteristics of all 54 individuals are summarized in **Table S1**, with human individuals who had received either the aP or wP vaccination during childhood being recruited for the study. All were eligible for TdaP (aP) booster vaccination. Study individuals were not recruited if they had pre-existing disease conditions (e.g., diabetes, hypertension).

#### Experimental data generation

Each multi-modal dataset consists of metadata about individuals and experimental data generated using five assays: plasma antibody measurements, PBMC cell frequencies, plasma cytokine concentrations, RNA sequencing, and T cell activation and polarization. We performed experiments on three pre-booster timepoints (days -30, -14, and 0) and five post-vaccine time points (days 1, 3, 7, 14, and 30).

1. <u>Plasma antigen-specific antibody measurements</u>. An indirect serological assay was employed using xMAP microspheres (Luminex Corporation) to measure Tdap antigen-specific antibody responses in human plasma. aP antigens (PT, PRN, Fim2/3, FHA), Tetanus Toxoid (TT), Diphtheria Toxoid (DT), and Ovalbumin (negative control) were coupled to uniquely coded beads (xMAP MagPlex Microspheres). A detailed description is provided by da Silva Antunes et al.^24^
2. <u>PBMC cell subset frequencies</u>. Twenty-one different PBMC cell subsets were measured by flow cytometry and manually gated using FlowJo (BD, version 10.7.0). A detailed description of is provided by da Silva Antunes et al.^24^
3. <u>Plasma cytokine concentrations</u>. Plasma samples were randomly distributed on 96-well plates for the quantification of different plasma cytokines by the Olink and/or Legendplex proteomics assays. A detailed description is provided by Willemsen et al. ^17^
4. <u>PBMC RNA sequencing</u>. Bulk RNA sequencing was performed on peripheral blood mononuclear cells (PBMCs) to measure transcriptional responses. Gene expression levels were quantified from paired-end sequencing reads and used for downstream analyses as described previously by da Silva Antunes et al.^24^
5. <u>CD4^+^ T-cell activation and polarization</u>. We performed Fluorospot (IL5/IFN-γ-producing cells) and activation by Fluorospot and AIM (the percentage of OX40^+^CD25^+^ CD4^+^ T cells) assays after aP antigen exposure. A detailed description is provided by Willemsen et al.^17^

Contestants were supplied with the baseline immunoprofiling data for all challenge dataset individuals. The post-vaccine response data, which contains the ground truth, was hidden from the contestants.

### Multi-step data harmonization pipeline

The training dataset comprised data from three independently collected cohorts (2020, 2021, and 2022). To enable robust modeling across years, we implemented a three-step preprocessing pipeline: 1) identifying common features, 2) baseline median normalization, and 3) batch-effect correction (**Figure 2A**).

As a first step, we identified shared features between the training cohorts and the 2023 challenge cohort to ensure consistency across datasets. Only features present in both were retained, resulting in 58,302 overlapping measurements (**Figure 2A**). Features are analytes measured in individual omics assays, such as cytokines in the plasma cytokine concentrations assay. Many of these features had low information content, especially for the transcriptomic assay. To address this, we filtered zero-variance and mitochondrial genes and removed lowly expressed genes (genes with transcript per million [TPM] < 1 in at least 30% of specimens) from the transcriptomic dataset. Similarly, we filtered features with zero variance from cytokine concentrations, cell frequency, and antibody assays. This resulted in 11,660 features: 11,589 transcriptomic, 28 cytokine, 20 antibody, and 23 PBMC cell frequency features.

In the second step, we performed assay-specific data normalization. We performed baseline normalization on cell frequency, antibody titer, and cytokine concentration data (**Figure S1**). Specifically, we calculated the median at baseline (days -30, -14, and 0) and used this value as a normalization factor. Each analyte value was divided by this factor. We did not apply any normalization to the gene expression data.

As a third step, we applied residual technical variability correction across the three training cohorts. We employed *ComBat,* an empirical Bayes method from the sva package, which models and removes batch effects while preserving biological variation^41,42^. Batch correction was applied separately within each assay. After batch-effect correction, we validated the effectiveness of this step by examining the distribution of features across batches (**Figures S2 and S3**). We noted a significant reduction in cross-year batch-associated variability, confirming that the batch-effect correction process was successful (**Figures S2 and S3**). This allowed us to move forward with a harmonized dataset for contestants for their analysis.

The challenge dataset was processed using the same data processing and normalization applied to the training dataset to ensure consistency and comparability (**Figure S1**). This included using the median of pre-vaccination data to normalize cell frequency, antibody titer, and cytokine concentration data. We did not apply any normalization to the gene expression data.

Both raw and processed versions were made available to contestants in TSV and RData formats, accompanied by the complete preprocessing code to support transparency and reproducibility.

### Formulating the prediction tasks

Contestants were challenged to predict a ranked list of the highest response (to be ranked first) to the lowest response (to be ranked last), and individuals for each prediction task were provided. We formulated six prediction tasks: three required contestants to predict specific biological readouts on particular days following the vaccine response, and the other three required contestants to predict the fold change between specific biological readouts on particular days following the vaccine response and the pre-vaccination state.

In Task 1.1, contestants were required to predict plasma IgG levels against PT on day 14 post-booster vaccination. For Task 1.2, contestants were required to predict the fold change of the plasma IgG levels against PT between day 14 post-booster vaccination and baseline. Tasks 2.1 and 2.2 required contestants to predict the overall frequency of monocytes among PBMCs on day 1 post-booster vaccination and the corresponding fold change, respectively. Similarly, Tasks 3.1 and 3.2 required contestants to predict *CCL3* gene expression on day 3 post-booster vaccination and the corresponding fold change values compared to baseline.

In addition to the main challenge, a separate bonus task (Task 4.1) was provided, which was independently evaluated, with a distinct task winner identified. Contestants were asked to predict and rank individuals based on their IFN-γ/IL-5 ratio (a surrogate for Th1/Th2 polarization) on day 30 post-booster vaccination. It is well established that aP-primed individuals have a polarization skewed towards Th2 responses (including IL-5 secretion), while wP-primed individuals exhibit a skew towards Th1 responses (including IFN-γ secretion)^17^. This divergence in T cell responses between aP and wP-primed individuals has been linked to differences in vaccine efficacy and durability and is maintained after booster vaccination^17^.

### Prediction challenge rules

1. Contestants were allowed to use both the provided training data and external data/predictors that could be used to train models.
2. Contestants were encouraged to draw inspiration from models and code used in the first and second challenges or apply similar techniques in this challenge. Contestants were welcome to build upon previous methods or utilize prior experience to make informed decisions. While there were no strict rules on reusing these models, we encouraged innovation rather than exact replication.
3. Contestants could submit as many submissions as they would like over the course of the challenge. The most recent submission by the submission deadline was counted as their final submission and was evaluated accordingly. It was possible to combine predictions from different models into one submission.
4. If contestants wanted to submit multiple final submissions, we asked that contestants to create a separate CMI-PB account for each submission. However, contestants had a maximum of 3 total final submissions.

### Performance evaluation metric

After receiving the contestants’ ranked predictions, we curated the rank file. *NA* values in the ranked list were imputed with the median rank for that list. Evaluations were then performed in two steps.

First, we chose the Spearman rank correlation coefficient as an evaluation metric to compare the predicted ranked list (*p*) for each task, *t*, and n individuals (n=21 for the set of challenge dataset individuals), *R_p,t_ = (r_p,1_, r_p,2_, …, r_p,n_)* against ground truth (*g*) ranked list *R_g,t_ = (r_g,1_, r_g,2_, …, r_g,n_)*. The Spearman rank correlation coefficient (ρ) is given by:

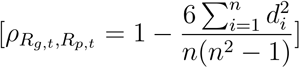

where *d_i_ = R_g, i_* - *R_p, i_* is the difference between the ranks of each pair. In this way, each task submitted by contestants was evaluated.

Second, we devised a point system to rank all submissions and identify the overall winner of the challenge. We awarded 3 points if a submission was top-ranked in a particular task and 1 point if the contestant attempted the task. The bonus task was evaluated separately only using the first step mentioned above, where winners were identified solely based on the Spearman rank correlation coefficient as an evaluation metric.

### Approaches for the top-three performing models

***1st Place Model: Random forest models with log-transformed data, PCA-reduced features, and tensor-based imputation.*** The team led by Cheng-Chang Wu from the University of Colorado first performed data preprocessing, wherein they extracted data from six time points: days 0, 1, 3, 7, 14, and 30 for the training set and day 0 for the challenge set (**Figure 3A**). Due to variation in features across years, only common features were retained for analysis. Prior to modeling, the assay data were normalized and standardized. A log(1 + x) transformation was applied to stabilize variance, followed by standardization to zero mean and unit variance. Within each data modality across years, features were filtered using two criteria: features with more than 50% missing values were removed, and features where more than 80% of observations were identical were excluded. Potential batch effects were addressed using the ComBat function from the SVA R package^43^, adjusting for infancy vaccine type and biological sex. To address missing data, all four data modalities and time points were combined into a tensor structure, and imputation was performed using a tensor decomposition-based method implemented via the tensorMiss R package^44^. This approach leveraged three-way correlations - across study individuals, features, and time points - to impute missing values. In addition to general imputation, features that exhibited identical values within specific batches (e.g., features “P60568” and “O95760” in the 2020 cohort) were treated as missing and similarly imputed.

Two modeling strategies were employed to predict post-vaccination immune responses: Random Forest and Generalized Additive Models with L1-penalized linear terms. Although imputation was performed on the entire dataset, model training was restricted to study individuals with observed outcome measurements, specifically IgG PT on day 14, monocytes on day 7, and *CCL3* on day 3. Predictor variables included clinical covariates such as infancy vaccine type, biological sex, and age of study individuals, the first 20 PCs from PCA, and the baseline measurement of the outcome of interest (e.g., baseline IgG PT for predicting day 14 IgG PT). For the Random Forest model, all predictors such as vaccine type, biological sex, age, baseline of the target outcome, and the first 20 PCs were included. In contrast, the Generalized Additive Model incorporated infancy vaccine type, biological sex, and the baseline of the target outcome as linear terms, each subjected to L1 regularization. Non-linear relationships were captured using penalized cubic splines with 10 knots for age, the PCs, and an interaction term between the first PC and vaccine type. To mitigate overfitting due to high degrees of freedom, the number of PCs included in this model ranged from one to five, with the optimal number selected via cross-validation.

Model performance was evaluated using 10-fold cross-validation, with mean squared error (MSE) used as the primary metric for model selection. The same trained model was used to generate predictions on both the original and fold-change scales. Fold-change predictions were derived by comparing model-predicted values with baseline (day 0) measurements.

***2nd Place Model: Cohort-aware modeling pipeline with nested cross-validation.*** The team led by Phillips Schaefer from Heidelberg University Hospital was interested in developing the most inclusive structured approach to account for the large-P-small-N problem^45^, multiple prediction tasks, diverse data modalities, and strong cohort effects. To address these, the workflow was designed around rigorous quality control, robust data preprocessing, and a carefully constructed model evaluation framework to minimize overfitting and ensure reproducibility (**Figure 3B**).

As the first step, the team developed a data normalization and integration approach to achieve assay-level consistency while systematically addressing cohort-specific peculiarities, such as differences in data units and quality warnings. For instance, inconsistent units in plasma antibody titers and cytokine concentration by Olink assays led to the exclusion of certain datasets to maintain data overlap between years. The 2020 cohort was excluded due to its underwhelming performance when integrated with later cohorts. Stringent quality control measures, such as excluding specimens with significant missing data or outliers, were implemented for assays like PBMC cell frequencies and plasma antibody titers. Various normalization techniques were employed, including median baseline normalization for PBMC cell frequencies and plasma antibody titers and variance-stabilizing transformations for PBMC gene expression. Integration strategies like ComBat-seq were used selectively for certain assays to address batch effects without overcorrecting^43^.

Model selection was guided by a nested cross-validation framework to ensure robust performance estimation under small-sample conditions. The outer loop employed group k-fold cross-validation to preserve cohort structure, while the inner loop used leave-one-out cross-validation (LOOCV) for hyperparameter tuning. A range of models, including LASSO, ElasticNet, and Random Forest, were evaluated based on predictive performance, variance across test folds, and compatibility with the available data. The team prioritized regularized models, and data from assays with frequent missingness, such as Olink, were excluded from predictive modeling to reduce noise. The team’s final pipeline demonstrated that refining data preprocessing and strategic model selection per task can yield reliable predictions, which the team suggested will provide a foundation for future applications of integrative immunological data analysis.

***3rd Place Model: XGBoost tree-based prediction of vaccine response metrics.*** The 3rd place team, led by Liran Mao from the University of Pennsylvania, developed a machine-learning approach using the XGBoost tree-based prediction method^33^, which used pre-vaccination data and early post-vaccination measurements (**Figure 3C**). Data preprocessing was critical in optimizing model performance. All assay datasets, except gene expression, were used in their preprocessed form supplied by the CMI-PB team. Gene expression data were normalized using a log_2_(TPM + 1) transformation to reduce skewness and stabilize variance.

Categorical variables, such as biological sex and infancy vaccination status (aP vs. wP), were one-hot encoded. Missing values were imputed using median imputation, and response variables were transformed to log2-fold changes to account for baseline variations. These data processing steps ensured that the data is well-prepared for modeling and helped maximize predictive accuracy while addressing challenges associated with small sample size and high feature dimensionality^45^.

Model selection involved benchmarking ten different machine learning algorithms, including linear (e.g., Lasso, Ridge, and Elastic Net) and non-linear models (e.g., Random Forest, Gradient Boosting, and XGBoost) (**Figure 3C**). Given the limited dataset size as compared to typical machine learning model building, performance evaluation used LOOCV, with root mean squared error (RMSE) as the primary loss function, and Pearson correlation was used for comparative performance assessment. XGBoost consistently outperformed other algorithms tested, particularly for absolute value tasks (1.1, 2.1, and 3.1). In contrast, prediction of fold-change tasks (1.2, 2.2, and 3.2) demonstrated varying model performance, suggesting distinct biological mechanisms influencing these task-specific responses. The final pipeline incorporated team-developed data preprocessing, hyperparameter-tuned XGBoost modeling, and prediction generation, complemented by rank-based transformations for challenge dataset samples.

### Quantification and statistical analysis

Statistical analyses are detailed for each specific technique in the specific Methods section or in the figure legends, where each specific comparison is presented. Statistical tests of the Spearman correlation coefficient were performed using R (version 4.1, www.r-project.org/) Details pertaining to significance are also noted in **Figure 2** legends, and p < 0.05 is defined as statistically significant.

## Data and code availability

The training and test datasets used for the first challenge can be accessible through our website at www.cmi-pb.org/. The web resource includes detailed information on the datasets, challenge tasks, submission format, submission files and evaluation code, descriptions, and access to the necessary data files that contestants used to develop their predictive models and make predictions. The codebase for standardizing data and generating computable matrices is available at www.cmi-pb.org/downloads/cmipb_challenge_datasets/current/3rd_challenge/. The codes for all models submitted for all three CMI-PB challenges are available, including those identified from the literature here on GitHub (https://github.com/topics/cmipb-challenge).

## Limitations of the study

The team by Liran Mao acknowledged that although XGBoost provided robust results, the analysis was constrained by the limited feature set. They suggested future directions could explore integrating additional immunological parameters and leveraging deep learning approaches to capture complex immune interactions contingent on larger datasets for improved generalization.

Commonly reported limitations included incomplete use of all available assay modalities, challenges in handling missingness at the assay level, and the risk of overfitting given the high-dimensional feature space and limited cohort sizes.

## Acknowledgements

We are grateful to the La Jolla Institute for Immunology’s flow cytometry, sequencing, and bioinformatics core facilities for their services. The authors would like to thank all study individuals who participated in the study and the clinical studies group staff, particularly Gina Levi, for all the invaluable help.

## Authors’ Information

### Contributions

P.S., S.O., M.K., and B.P. organized the CMI-PB Challenge. The top-performing three approaches were designed by the following teams: A.G., C.C.W., Liran M., C.L., Y.T., T.A.N., N.S.C., and P.S.L.S. The remaining approaches were proposed by CMI-PB Challenge contestants. P.S., B.P. interpreted the results of the challenge and all follow-up analyses. P.S., B.P. wrote the paper with input from all other authors. J.L. and L.W. generated the experimental data. T.E., L.G., F.A., B.G., S.H.K., and B.P. supervised the research.

