## Supplementary Materials for "An Open Benchmark for Systems Vaccinology: Insights from the CMI-PB Challenges"


#### Supplementary Tables

**Table S1. The characteristics of all 54 subjects in the challenge dataset.** For each subject, the table lists the subject identifier, age in years at enrollment, biological sex at birth, and childhood pertussis vaccine priming status (aP, acellular; wP, whole-cell). The cohort spans ages 19 to 36 years (median 26) and comprises 33 female and 21 male subjects, split evenly between aP (n=27) and wP (n=27) priming groups.

| SubjectID | Age | BiologicalSexAtBirth | VaccinePrimingStatus |
|---|---|---|---|
| 119 | 23 | Female | aP |
| 120 | 27 | Female | wP |
| 121 | 22 | Female | aP |
| 122 | 23 | Female | aP |
| 123 | 26 | Female | wP |
| 124 | 22 | Male | aP |
| 125 | 29 | Male | wP |
| 126 | 29 | Male | wP |
| 127 | 26 | Female | aP |
| 128 | 28 | Female | wP |
| 129 | 31 | Male | wP |

| | | | |
|---|---|---|---|
| 130 | 26 | Male | wP |
| 131 | 24 | Female | aP |
| 132 | 27 | Male | wP |
| 133 | 25 | Female | aP |
| 134 | 32 | Male | wP |
| 135 | 27 | Male | wP |
| 136 | 27 | Female | wP |
| 137 | 24 | Female | aP |
| 138 | 22 | Male | aP |
| 139 | 29 | Female | wP |
| 140 | 21 | Female | aP |
| 141 | 26 | Female | wP |
| 142 | 31 | Female | aP |
| 143 | 19 | Female | aP |
| 144 | 23 | Female | aP |
| 145 | 20 | Male | aP |
| 146 | 31 | Male | wP |
| 147 | 23 | Female | aP |
| 148 | 35 | Male | wP |
| 149 | 32 | Female | wP |
| 150 | 32 | Male | wP |
| 151 | 31 | Female | wP |
| 152 | 28 | Female | wP |
| 153 | 25 | Female | aP |
| 154 | 26 | Female | aP |
| 155 | 26 | Female | aP |

| | | | |
|---|---|---|---|
| 156 | 22 | Female | aP |
| 157 | 26 | Female | aP |
| 158 | 23 | Male | aP |
| 159 | 29 | Female | wP |
| 160 | 27 | Male | aP |
| 161 | 30 | Female | aP |
| 162 | 24 | Female | aP |
| 163 | 30 | Female | wP |
| 164 | 32 | Male | wP |
| 165 | 30 | Female | wP |
| 166 | 22 | Female | aP |
| 167 | 26 | Male | aP |
| 168 | 32 | Male | wP |
| 169 | 20 | Male | aP |
| 170 | 31 | Male | wP |
| 171 | 20 | Female | wP |
| 172 | 36 | Male | wP |

**Table S2. Antibody panel used for flow cytometry.** The table lists the 20 antibodies used for surface staining, giving the target antigen, fluorochrome conjugate, host species, target species, clone, vendor catalog number, vendor, and working dilution.

| Target | Conjugate | Host | Target | Clone | Catalog | Vendor | Dilution |
|---|---|---|---|---|---|---|---|
| CD45 | BUV395 | Mouse | Human | HI30 | 563792 | BD | 1/500 |
| CD8 | BUV496 | Mouse | Human | RPA-T8 | 612942 | BD | 1/500 |
| CD20 | BUV563 | Mouse | Human | 2H7 | 748456 | BD | 1/200 |
| CD16 | BV510 | Mouse | Human | 3G8 | 612786 | BD | 1/100 |
| CD3 | BUV805 | Mouse | Human | UCHT1 | 612895 | BD | 1/200 |
| CD14 | BV480 | Mouse | Human | M5E2 | 746304 | BD | 1/100 |
| CD45RA | BV570 | Mouse | Human | HI100 | 304132 | Biolegend | 1/200 |
| CD19 | BV605 | Mouse | Human | HIB19 | 302244 | Biolegend | 1/200 |
| IgD | PE-CF594 | Mouse | Human | IA6-2 | 747484 | BD | 1/200 |
| CD11c | BV785 | Mouse | Human | 3.9 | 301644 | Biolegend | 1/50 |
| CCR7 | FITC | Mouse | Human | G043H7 | 353216 | Biolegend | 1/66 |
| CD123 | PE-Cy7 | Mouse | Human | 6H6 | 306016 | Biolegend | 1/100 |
| CD38 | PerCP-Cy5.5 | Mouse | Human | HIT2 | 562288 | BD | 1/50 |
| HLA-DR | AF700 | Mouse | Human | L243 | 307616 | Biolegend | 1/50 |
| CD56 | APC | Mouse | Human | 5.1H11 | 362504 | Biolegend | 1/100 |
| CD4 | APC-eF780 | Mouse | Human | RPA-T4 | 47-0049-42 | LIFE TECH | 1/200, 1/50 |
| CD71 | PE-Cy5 | Mouse | Human | M-A712 | 551143 | BD | 1/50 |
| CD66b | BV421 | Mouse | Human | G10F5 | 562940 | BD | 1/100 |
| CD1c | BV650 | Mouse | Human | 1.161 | 331542 | Biolegend | 1/200 |
| CD141 | PE | Mouse | Human | M80 | 47-0049-42 | LIFE TECH | 1/200 |

### Supplementary Figures

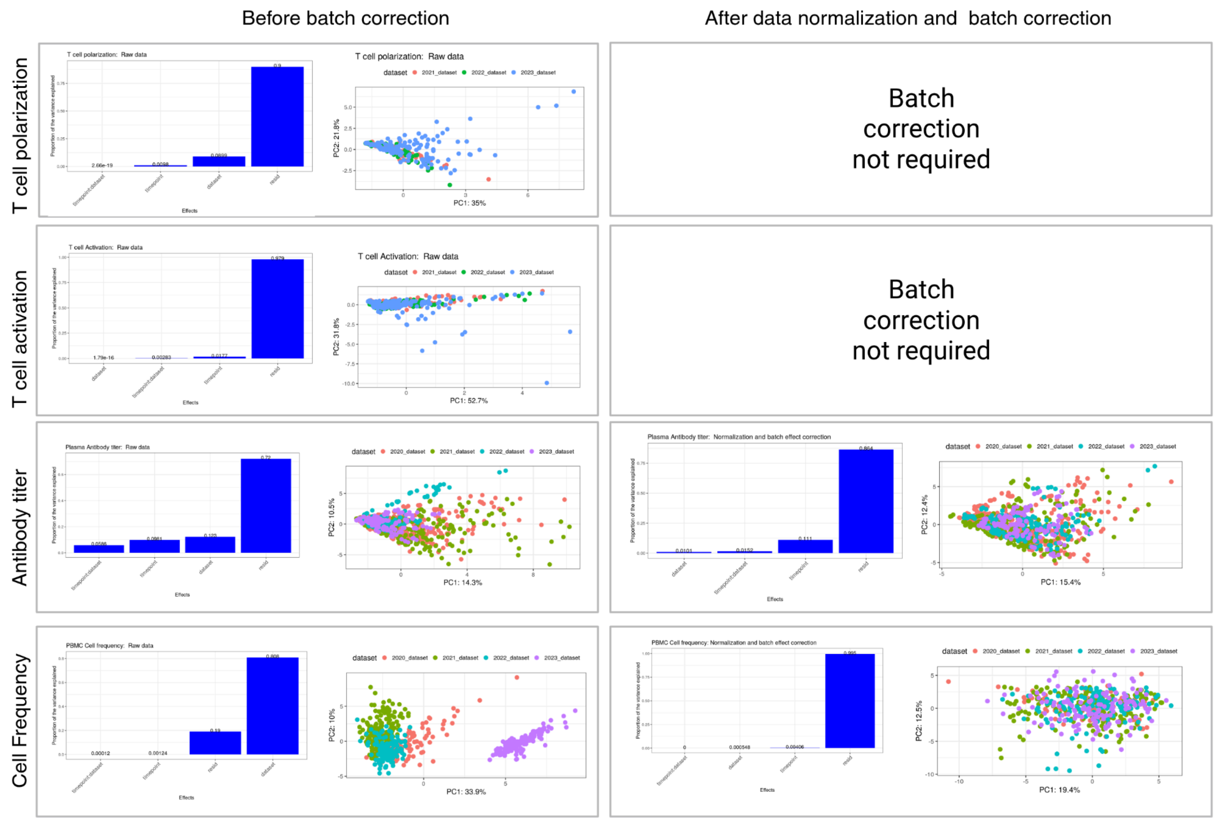


**Figure S1: Plot of assay data before and after normalization and batch effect correction for T cell apolarization, T cell activation, ab titer, and Cell frequency data.** For each assay, the left plots show data before batch correction, while the right plots show data after normalization and batch correction.

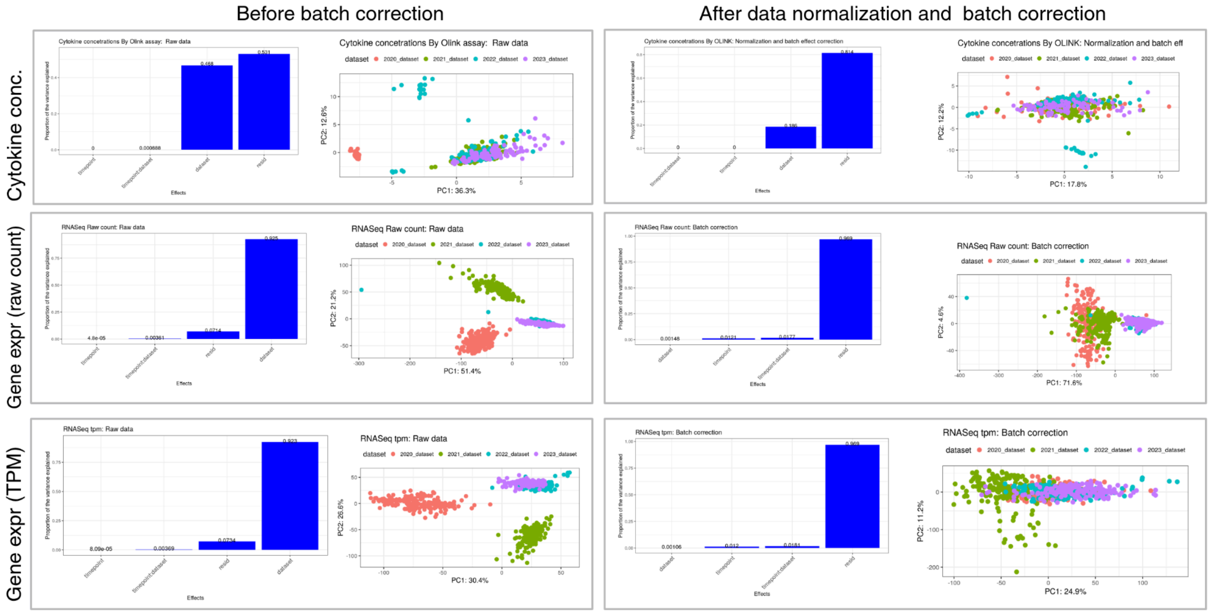


**Figure S2: Plot of assay data before and after normalization and batch effect correction for Gene Expression and Cytokine Concentration data.** For each assay, the left plots show data before batch correction, while the right plots show data after normalization and batch correction.

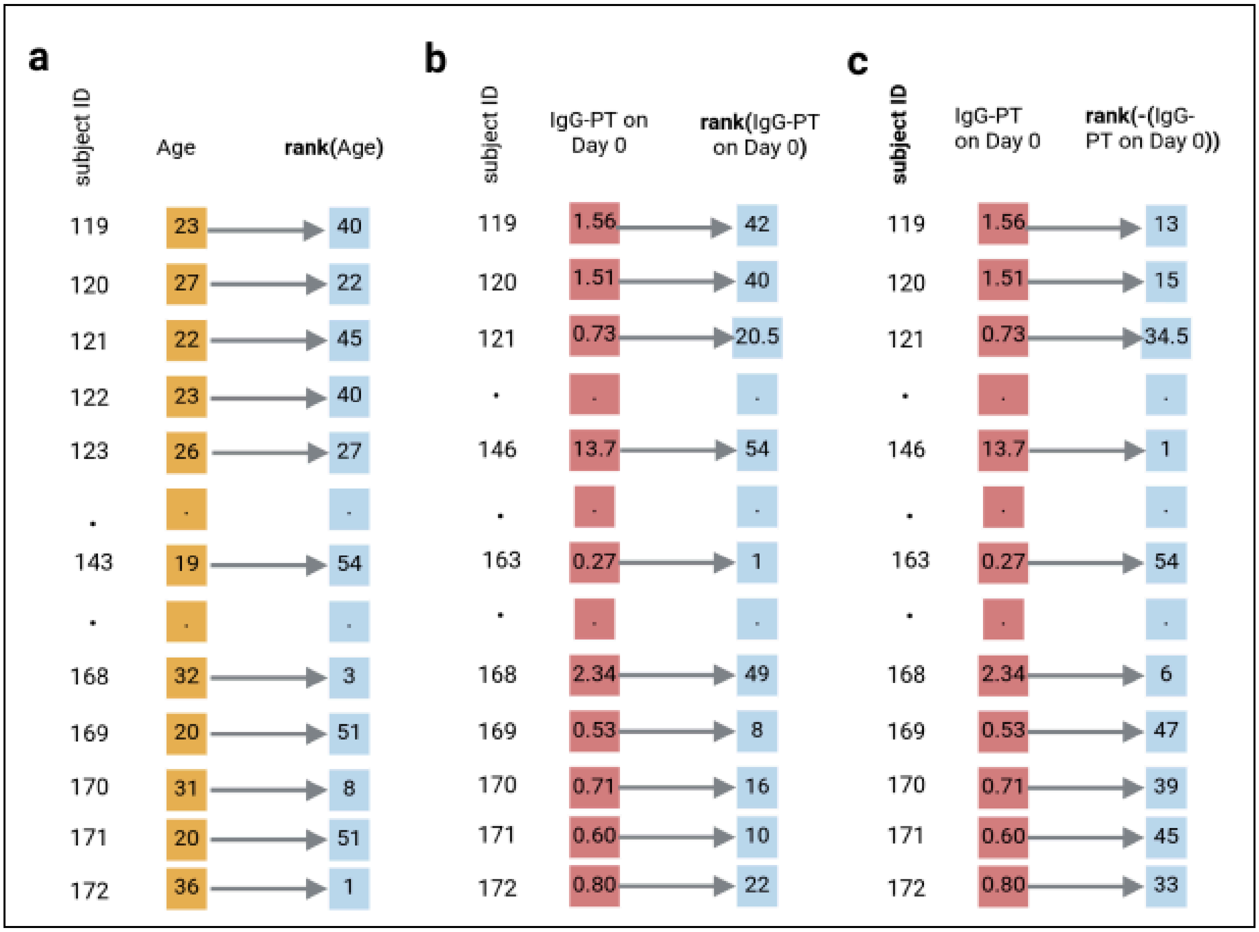


**Figure S3: Schematic of baseline control model construction across tasks.** (a) Demographic control: Raw age values (orange) are ranked (blue) to generate a subject-level ranking based on age alone. (b) Absolute-level tasks (Tasks 1.1, 2.1, 3.1): Baseline measurements (e.g., IgG-PT at day 0, monocyte frequency at day 1, CCL3 expression at day 3; red) are ranked (blue) to reflect subject ordering by pre-vaccination immune status. (c) Fold-change tasks (Tasks 1.2, 2.2, 3.2): The inverse of baseline measurements (red) is used prior to ranking (blue) to prioritize individuals with lower baseline values, simulating the potential for greater fold-change responses post-vaccination.

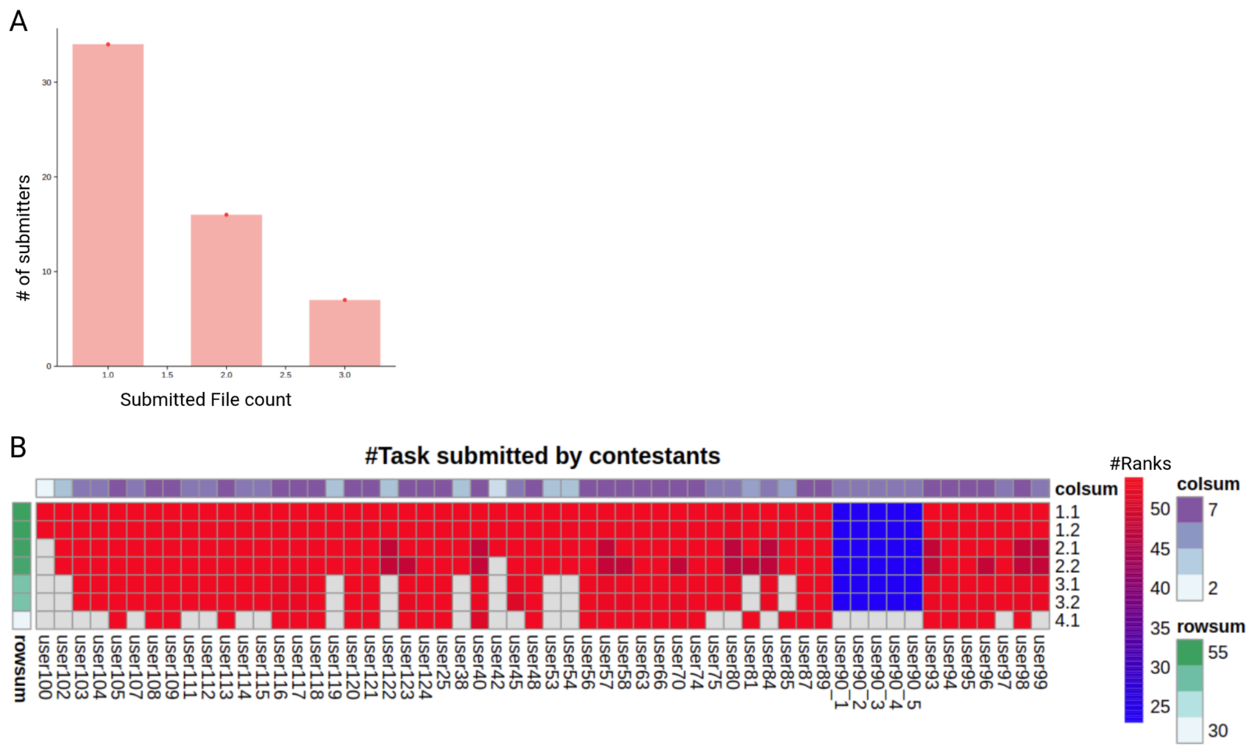


**Figure S4. Contestant submissions and task coverage in the public CMI-PB challenge.** (A) Distribution of the number of final submissions per team, showing the count of teams that submitted one, two, or three models. (B) Heatmap summarizing task participation across teams. Rows represent prediction tasks (1.1-4.1), and columns represent individual teams. Colored cells indicate tasks attempted by each team. Sidebars denote the total number of tasks attempted per team (column sum) and the number of teams attempting each task (row sum).

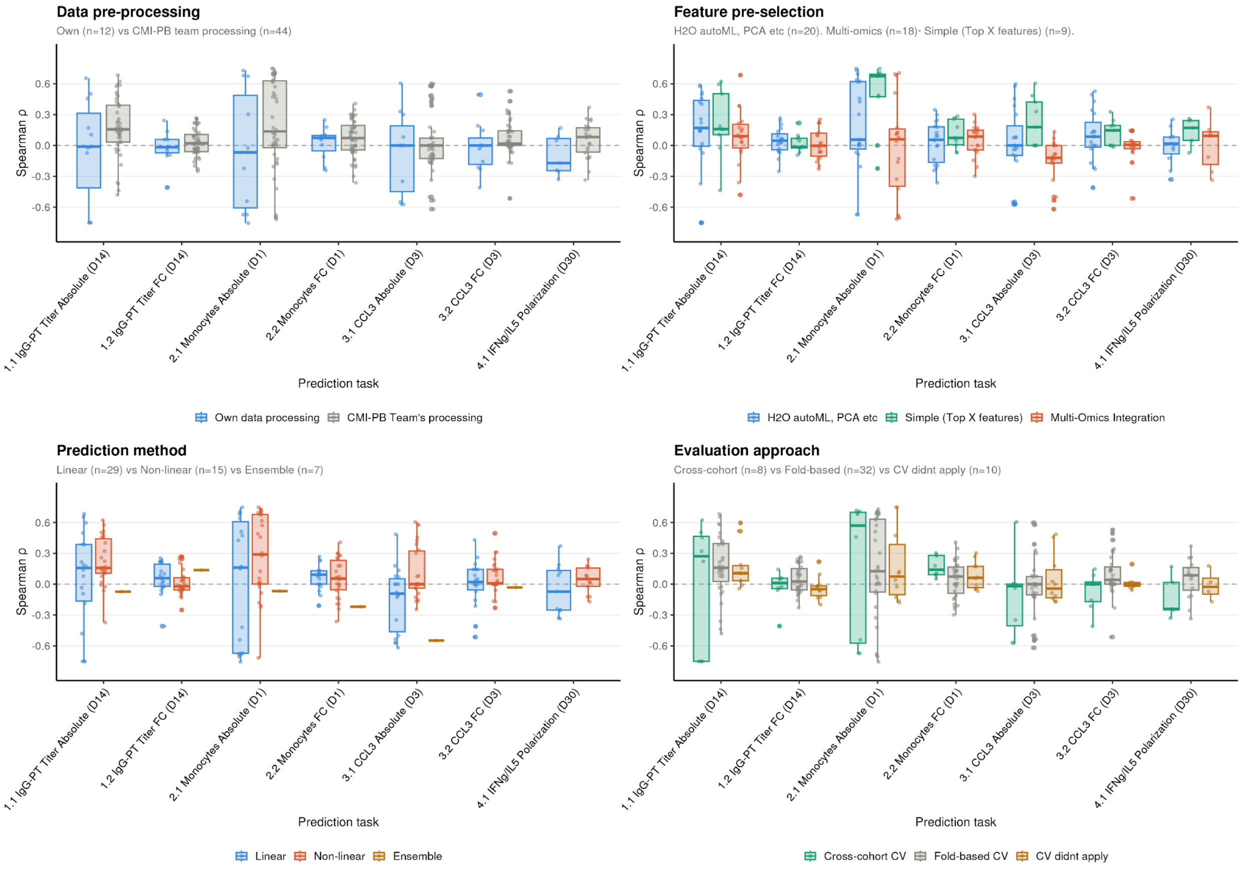


**Figure S5. Model performance across computational modeling strategy categories in the CMI-PB third challenge.** Boxplots show the distribution of Spearman correlation coefficients (ρ) between predicted and observed post-vaccination immune responses for 54 submitted models across seven prediction tasks. Each panel compares model performance grouped by a distinct methodological choice: (top left) data pre-processing approach (own processing, n=12; CMI-PB team processing, n=44); (top right) feature pre-selection strategy (H2O autoML/PCA, n=20; simple top-X features, n=9; multi-omics integration, n=18); (bottom left) prediction method (linear, n=29; non-linear, n=15; ensemble, n=7); (bottom right) cross-validation approach (cross-cohort CV, n=8; fold-based CV, n=32; CV not applied, n=10). Individual points represent single model–task pairs. The dashed horizontal line marks ρ = 0. Box boundaries indicate the interquartile range (IQR), the center line the median, and whiskers extend to 1.5×IQR. Tasks are shown along the x-axis: absolute and fold-change readouts for IgG-PT titer (Day 14), monocyte frequency (Day 1), CCL3 gene expression (Day 3), and IFNγ/IL-5 Th1/Th2 polarization ratio (Day 30).

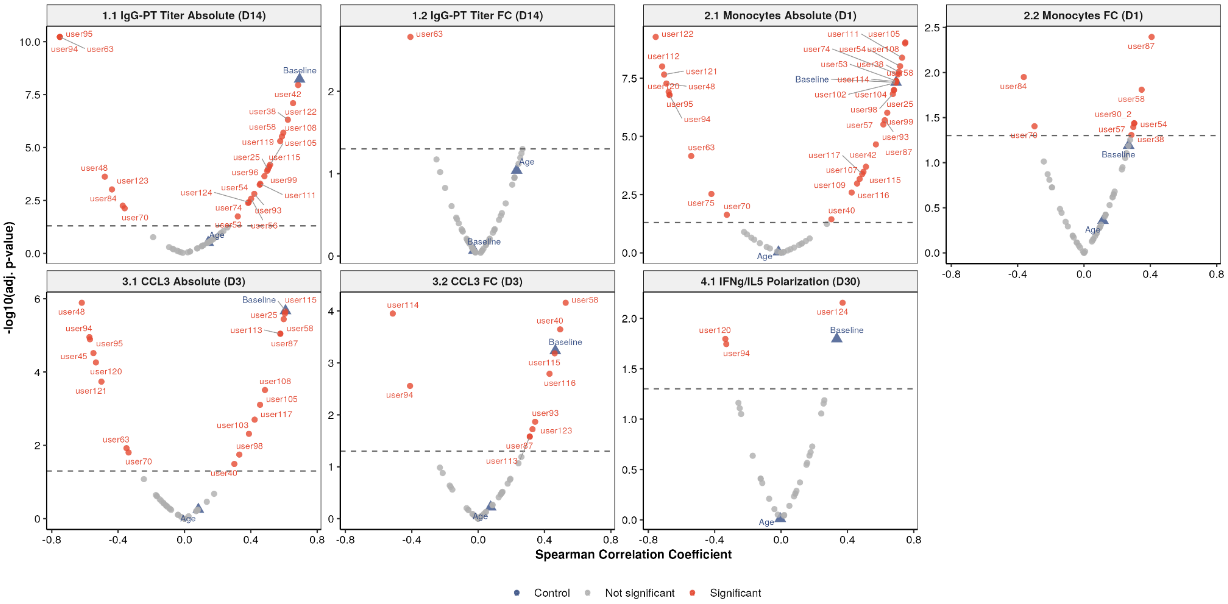


**Figure S6. Spearman correlation analysis between contestant-submitted predictions and measured post-vaccination outcomes across CMI-PB challenge tasks.** Volcano-style plots show Spearman correlation coefficients (x-axis) versus $-\log_{10}$ adjusted p-values (y-axis) for baseline features correlated with IgG-PT titer absolute and fold-change (FC) at Day 14 (1.1–1.2), monocyte absolute and FC at Day 1 (2.1–2.2), CCL3 absolute and FC at Day 3 (3.1–3.2), and IFNγ/IL-5 polarization at Day 30 (4.1). Red circles indicate significant correlations (FDR-adjusted $p < 0.05$, dashed line); gray circles are non-significant. Blue triangles denote reference variables (baseline, age).

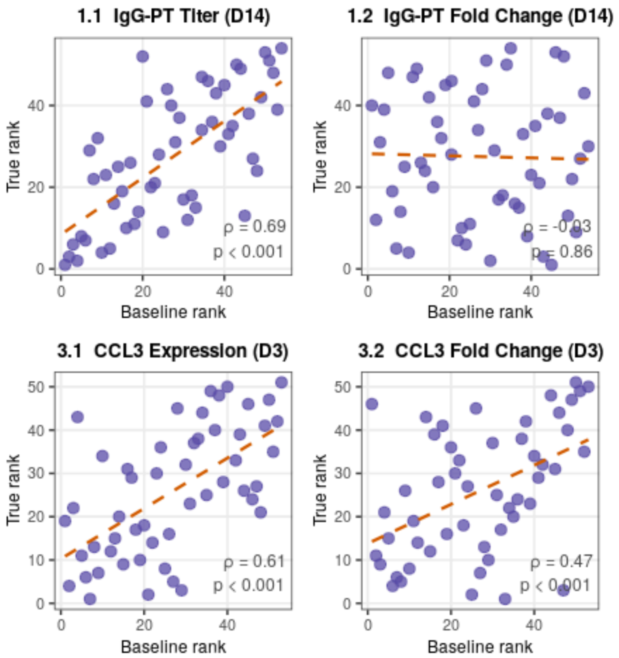


**Figure S7. Pre- versus post-vaccination immune readouts across 54 participants.** Scatter plots show pre-vaccination (x-axis) versus post-vaccination (y-axis) values for IgG-PT antibody titers (Day 14), monocyte frequencies (Day 1), and CCL3 gene expression (Day 3). Each point represents one individual. The Spearman correlation coefficient and adjusted p-value are indicated per panel. Regression line shown in gray. The spread of post-vaccination values among individuals with similar pre-vaccination levels illustrates the degree of inter-individual variability that remained unexplained by baseline state alone.
